# Microbial community structure and ecology in sediments of a pristine mangrove forest

**DOI:** 10.1101/833814

**Authors:** C.O. Santana, P. Spealman, V.M.M Melo, D. Gresham, T.B. Jesus, F.A. Chinalia

## Abstract

Mangrove forests are coastal intertidal ecosystems, characterized by mangrove trees growing in slow moving saline waters, that constitute a large portion of the coastline in the tropical and subtropical regions. The dynamic water regime created by the tides results in different microhabitats in which microbial communities play an essential role in the functioning and maintenance of the mangrove ecosystem. However, little is known about the diversity of taxa within these micro-habitats and their functional roles, as only a small fraction of these organisms can be cultured in the laboratory. In this study, we characterized the microbial community present in three distinct regions of mangrove sediments from the Serinhaém estuary, part of the Atlantic Forest biome within the Environmental Protection Area of Pratigi. We sampled sediments from regions below the tidal waterline (submerged), intertidal regions (intertidal), and regions above the tidal waterline (seco). More than 85% of all the sequences in the samples belonged to 6 of 42 identified phyla: *Proteobacteria* (30.6%), *Firmicutes* (30%), *Chloroflexi* (8.7%), *Planctomycetes* (5.7%), *Crenarchaeota* (5.4%) and *Actinobacteria* (5.3%). Diversity indices show that the submerged regions of the mangrove forest exhibit the greatest diversity and richness relative to the other regions. Notably, the intertidal region has the least diversity, suggesting that the dynamics of environmental variables in this region has an important influence on microbial diversity. Furthermore, distance metrics indicate that submerged sediments are more homogeneous while the seco region exhibits greater variability between locations. Finally, we found that the most abundant microbial families in the sediments are associated with nutrient cycling consistent with the essential role of the microbiome in maintaining the health of the mangrove ecology.

## INTRODUCTION

Soils are among the greatest sources of microbial diversity on the planet (Tveit et al. 2013; Kaur et al. 2015), are fundamental to terrestrial processes such as carbon sequestration and nitrogen cycling, and can shape important characteristics of habitats through metabolic activities (Wendt-Potthoff et al. 2012). However, little is known about the microbial diversity and functional roles exerted by different taxa as only a small fraction of these organisms can be cultured in the laboratory (Mocali and Benedetti 2010; Kaur et al. 2015; Bornemann et al. 2015). Thus, metagenomic approaches are needed to establish a basis for assessing community changes in response to environmental disturbances or anthropogenic pollution (Mahmoudi et al. 2015).

Mangrove ecosystems constitute a large portion of the coastline in the tropical and subtropical regions of Earth and are characterized by their salinity and tidal variation which results in frequent anaerobic conditions and a wide range of redox potential. Such conditions make mangroves hotspots for microbial diversity, and the microbial community plays essential roles in the functioning and maintenance of the ecosystem (Andreote et al. 2012).

The Atlantic Forest in Brazil is recognized as one of the most biodiverse ecosystems on the planet, containing mangroves, forests and restinga fields but is threatened by anthropogenic disturbances such as logging and farming resulting in a severe decline in its original area (Ditt et al. 2013; Brasil, 2010). However in the south part of Bahia State, Brazil, it is still possible to find a territorial band of great environmental relevance with significant fragments of the Atlantic Forest In this region is located in the Environmental Protection Area (APA) of Pratigi, created in 1998 with the aim to preserve this important Atlantic rainforest fragment (MMA 2004). Recent studies evaluating the environmental conditions in the APA have shown that this preservation effort has been effective (Lopes et al. 2011); (Ditt et al. 2013); (Mascarenhas et al. 2019; “Website” n.d.)). However, the microbial communities within the mangrove forests of the APA remain poorly defined.

In this study we characterized the prokaryotic microbiota present in mangrove sediments of the Serinhaém estuary. We assessed the structure of microbial communities, the influence of environmental variables on the diversity of these communities, and identified possible drivers for the different nutrient cycles. Our study provides insight into the role of microbes in the functioning of mangrove forests and establishes a framework for monitoring the health of this important ecosystem.

## METHODS

### Study area

The Serinhaém Estuary is located in the Low South Region of Bahia State, Brazil (Fig. 1), between the coordinates 13°35’N and 14°10’S and 39°40’W and 38°50’E. The estuary is within the Pratigi Environmental Protection Area (APA), one of the remaining Atlantic forest region with a total area of 85,686 ha, enclosing a 32 km long portion of the lower Juliana River and emptying directly into Camamu Bay along with several smaller rivers (Corrêa-Gomes et al. 2005).

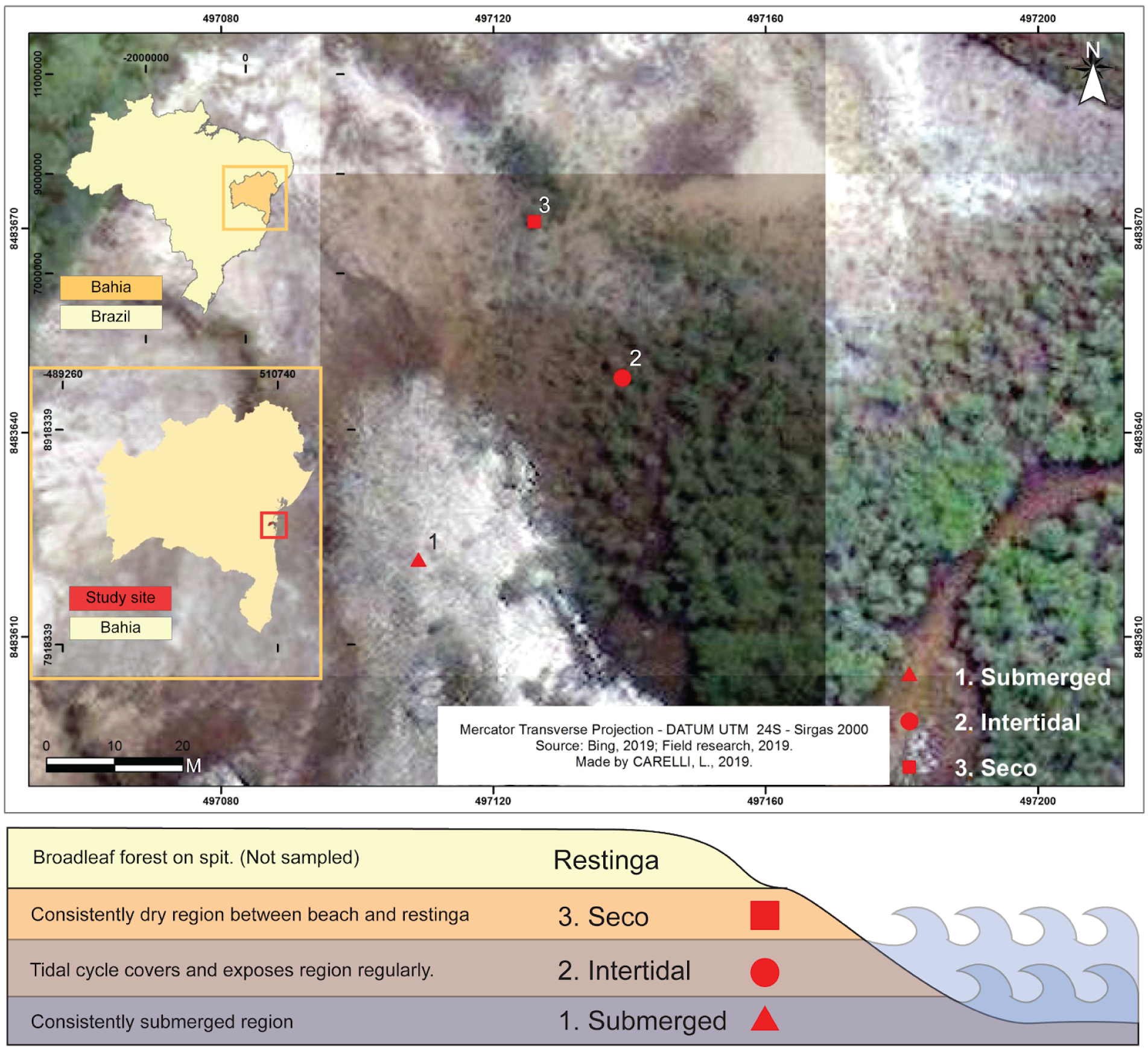

### Sampling and DNA extraction

Samples were collected from 3 sites on the Serinhaém estuary in July 2018. Physical-chemical parameters such as temperature, salinity and dissolved oxygen in the water column were measured using a multiparameter monitoring system (YSI model 85, Columbus). Sediment samples were collected in triplicate at each site with a sediment core (10 cm of the surface layer) and transferred to the Laboratory of Petroleum Studies (LEPETRO) at the Federal University of Bahia (UFBA). Submerged, Intertidal, and Seco regions were sampled (Supplemental Figure 1). For each sediment core an aliquot was separated and frozen for subsequent DNA extraction while the remainder of the sample was used for measuring the organic matter content. The total genomic DNA was extracted from 0.25 g of soil using the Power Soil DNA Isolation Kit (Qiagen, Carlsbad, CA, USA) in the Microbial Ecology and Biotechnology Laboratory (LEMBiotec) at the Federal University of Ceará (UFC). All DNA samples were stored at −20° C before analysis.

### Library preparation and sequencing

After DNA extraction, we used PCR to specifically targeting the V4 region of the bacterial 16S rRNA with the primer-pair 515F-Y (Parada, Needham, and Fuhrman 2016) and 806R-XT (Caporaso et al. 2011). Sequencing of the DNA present in the sediment samples was performed through the Illumina MiSeq platform using the V2 kit (300 cycles). Following demultiplexing the data were stored in the BaseSpace platform for subsequent bioinformatic analysis.

### Data analysis

Trimmomatic was used to quality filter and trim demultiplexed sequences (ILLUMINACLIP:TruSeq3-PE.fa:2:30:10 LEADING:3 TRAILING:3 SLIDINGWINDOW:4:15 MINLEN:100). Trimmomatic parameters were defined using FastQC. After trimming forward and reverse reads were joined into single reads using QIIME1 (join_paired_ends.py, -j 4 -p 1). This resulted in reads of approximately 250 bp in length which were used for all subsequent steps. Reads were denoised using DADA2 (denoise-single, --p-trim-left 3, --p-trunc-len 0) in QIIME2 (q2cli, version 2019.4.0). Denoised sequences were separated into Operational Taxonomic Units (OTUs), which can be defined as clusters of similar sequence variants of the same gene region. The OTU picking step was performed with the Vsearch tool in QIIME2 using the Open Reference method with a 97% similarity (--p-perc-identity 0.97) against the reference 16S rRNA sequences in Greengenes database (version gg_12_10). Phylogenetic reconstruction was performed in QIIME2 resulting in alignment and phylogenetic tree files.

Tree visualization was performed with R (version 3.4.4) using the Metacoder package (version 0.3.2). Core metric analyses and group significance tests were carried out for alpha and beta diversity in QIIME2. Posterior analysis was performed using the R Phyloseq package (version 1.22.3). Posterior analyses were plotted in in R using ggplot2.

Correlations between community structure and environmental variables were tested and displayed using the Vegan package in R (version 2.5-6). Taxonomy assignment was carried out in QIIME2 using the representative sequences for each OTU and the Greengenes classifier for V4 16S rRNA gene region, thereby allowing the identification of microorganisms groups in the samples. Groups that accounted for more than 3 sequences and were present in at least 2 samples from the data set were chosen as representatives for correlation analysis in order to remove the taxa with very low frequencies. The QIIME2 zip files generated by the pipeline were exported and transformed in R, with the QIIME2R package (0.99.12 version).

Identification of site specific taxa enrichment was performed by first normalizing replicates by downsampling OTU abundance to match the least abundant replicate (Submerged replicate 2). The means of normalized observations were used to determine if differences in OTU abundances between sites were significant using a chi-squared test. To account for differences could be due to variance between replicates we further required that the distributions of unnormalized OTU abundances between sites be significantly different using a Mann-Whitney U test (p-val <= 0.05). Finally, because higher abundant taxa can have statistical significance with minimal biological significance we also required the effect size to be in excess of a 5% difference in log-fold abundance between sites.

The entire computational workflow is available as a repository in github: https://github.com/GreshamLab/Santana-et-al-2019.

All raw data are available on OSF: https://osf.io/h6k5s/

## RESULTS

### Taxonomic composition of prokaryotic communities

After quality filtering, a total of 204,599 bacterial and archaeal sequences remained for community analysis, corresponding to an average of 22,733.2 sequences per sample. Sequence clustering yielded a total of 1,708 OTUs. Of these, 1,607 OTUs and 193,153 sequences were assigned to Bacteria (94.1%) and 101 OTUs and 114,44 sequences were assigned to Archaea (5.9%) kingdoms. Only 2 sequences could not be assigned to any prokaryotic kingdom. From all the identified OTUs it was possible to differentiate 42 unique phyla, 117 classes, 136 families, 96 genera and 39 species. More than 85% of all the sequences in the samples belonged to 6 of the 42 phyla: *Proteobacteria* (30.6% abundance, 61,889 sequences), *Firmicutes* (30% abundance, 60,618 sequences), *Chloroflexi* (8.7% abundance, 61,889 sequences), *Planctomycetes* (5.7% abundance, 11,592 sequences), *Crenarchaeota* (5.4% abundance, 10964 sequences) and *Actinobacteria* (5.3% abundance, 10853 sequences). The total sequence and OTU abundances in all the observed phyla is summarized in Supplemental Table 1. Figure 2 shows all the classes, orders and families belonging to the 6 dominant phyla in the data set. We also quantified taxa for each site separately (Supplemental figures 2, 3 and 4).

**Figure 2.**
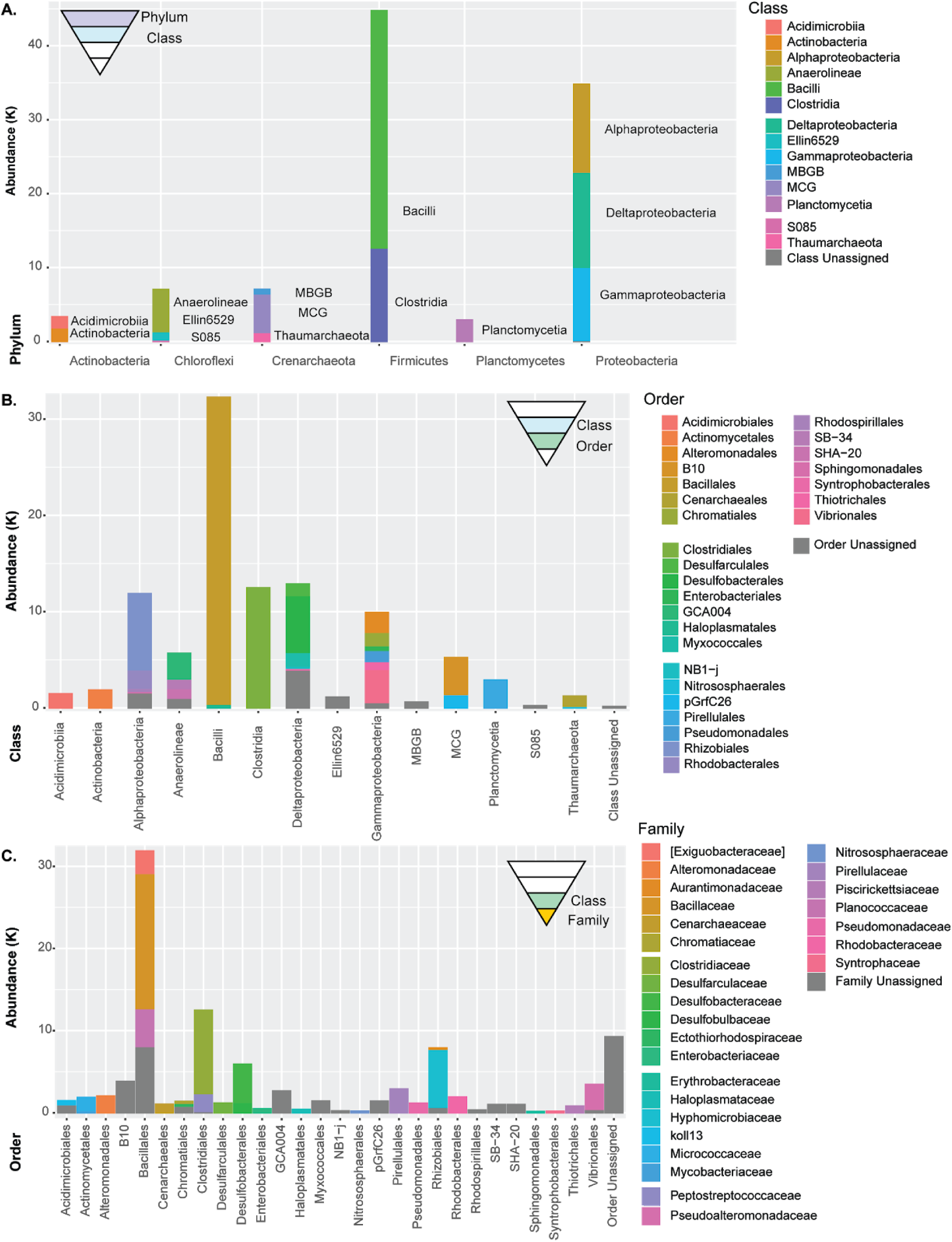
Taxonomic abundances from all sample sites. *Proteobacteria* (30.6% of total) and *Firmicutes* (30%) make up the majority of the phyla. Considered with *Chloroflexi* (8.7%), *Planctomycetes* (5.7%), *Crenarchaeota* (5.4%), and *Actinobacteria* (5.3%) account for more than 85% of all taxa identified. For this presentation purposes we filtered taxa with abundances less than 1% of the total.

**Figure 3.**
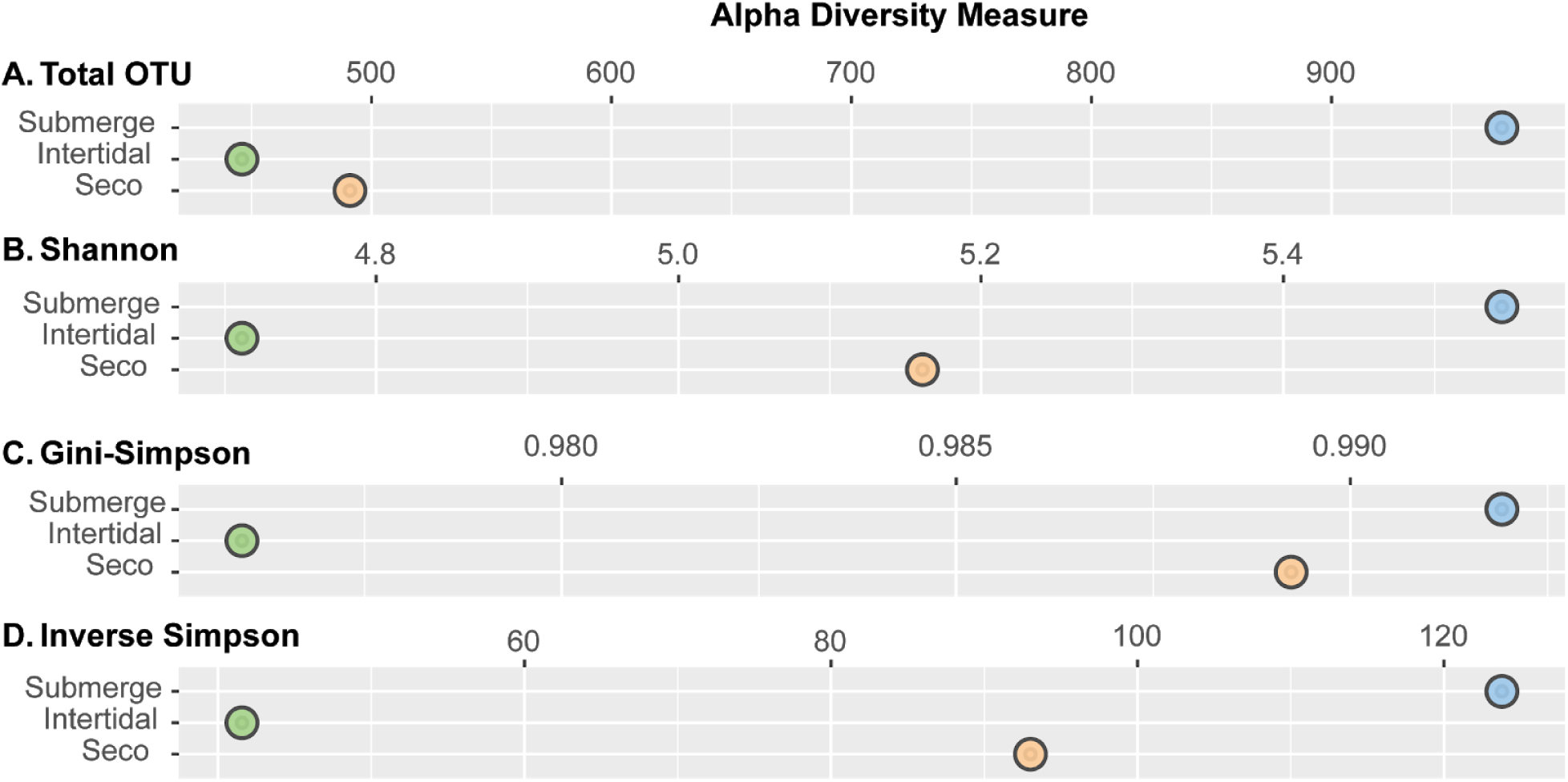
Alpha diversity by sampling site. Alpha diversity (mean diversity of species per site) measured using the absolute number of OTUs (**A**), taxa richness and abundance normalized measures (**B, C**). A measure of the ‘true diversity’ species number equivalents (**D**).

**Figure 4.**
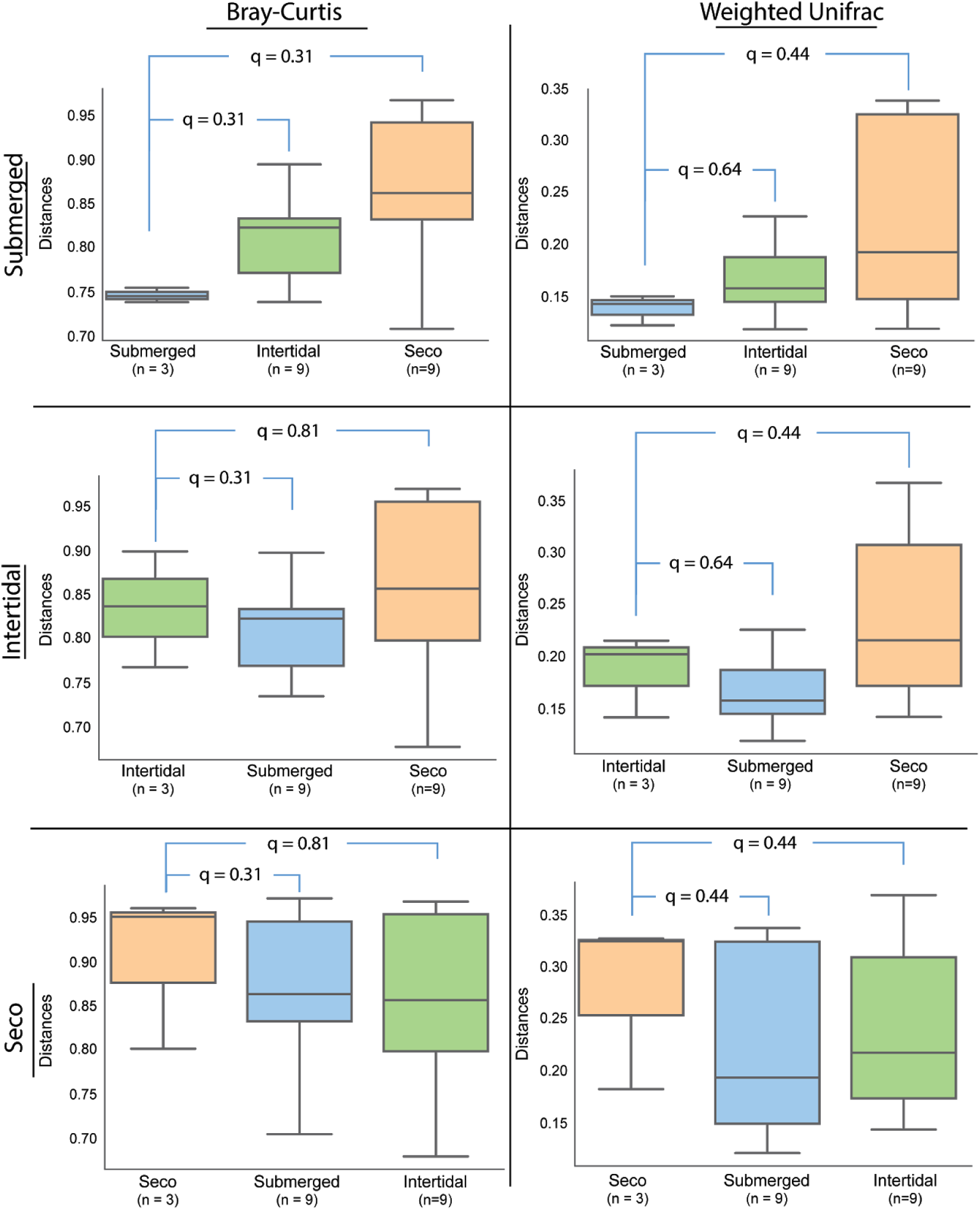
Beta-diversity between sampling sites. Beta-diversity is the measure of taxonomic difference between any two sites. All sites are compared pairwise both Bray-Curtis and Weighted Unifrac quantitations. We find strong agreement between both methods.

### Alpha diversity by site shows communities at Submerged sites to be the most diverse

The alpha diversity indices for each of the sampling sites are shown in Figure 3. Overall the sediments from the submerged mangrove sites had the greatest values for richness and diversity indices, while the intertidal region had the lowest alpha diversity indices.

The Total OTU accounts for the richness in each site and is the absolute number of OTUs found in the samples, not considering abundances (Fig 3A). We find the number of OTUs in the submerged region of the mangrove to be nearly twice as large as the number found for the seco region and even greater compared to the richness estimates for the intertidal samples. In addition to the Alpha-diversity metrics described above, we also used additional estimators Chao1 and ACE (Supplemental Figure 5), which show the same general trend.

**Figure 5.**
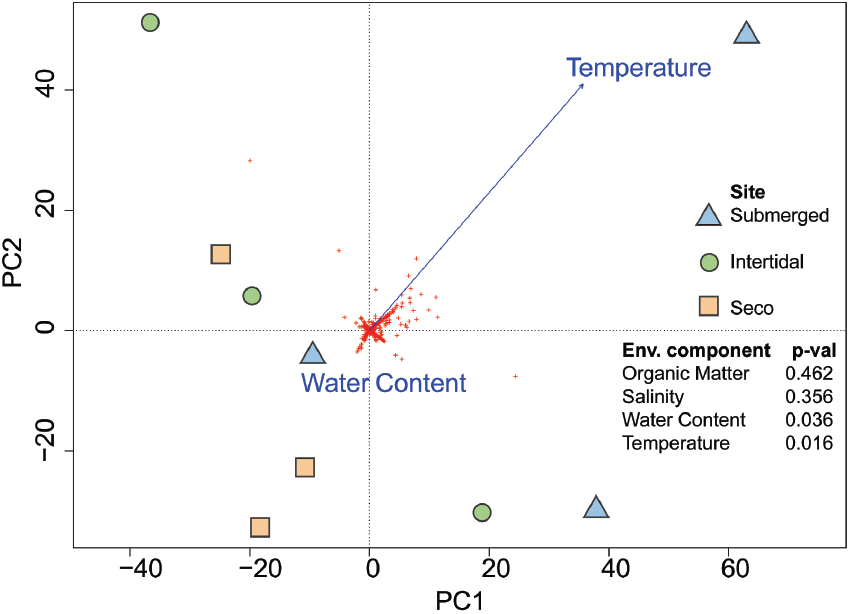
Principal Component Analysis of environmental variables finds Temperature and Water Content to be significant correlates with prokaryotic communities. The plot is displaying the two significant environmental variables that influence the prokaryotic communities in the samples and the p-values (as calculated by vegan from random permutations of the data, eg. Pr(>r)) for all tested correlations. Water content clusters in the x axis as it is a factor variable and temperature is displayed as a vector in which length (importance of the influence) and direction can be observed.

Diversity indices are mathematical measures based on taxa richness and abundance. Shannon’s diversity index ranges from a high of 5.54 in the submerged regions to 4.71 in the intertidal samples (Fig 3B). However, because diversity indexes usually involve non-linear scales with different sensitivities to rare and common species, the index can be a nonintuitive measure. By transforming diversity indices into true diversities (Jost 2006) we can explore the information using the more intuitive measure of number equivalents (Jost 2007). The Shannon’s true diversity for the submerged site means that this site has the same diversity as a site having 245 species evenly abundant while the intertidal site is as diverse as having 110 equally frequent species and the seco site has a diversity equivalent to 181 equally abundant species. This suggests that the submerged regions are more than 50% greater in diversity than the intertidal region that experiences great variations in water regimen and at least 25% more diverse than the seco site of the mangrove forest.

The Gini-Simpson’s diversity index is more sensitive to OTU abundance than Shannon’s diversity, giving more weight to the most frequent OTUs in the samples (Fig 3C). This index also indicates higher diversity in the sediments from the submerged sites compared to the intertidal region. The Gini-Simpson’s index can also be represented by true diversities (Fig 3D). The diversity shown by submerged sediments is equivalent to 124 equally abundant species while the diversity in the seco and intertidal area sediments are 93 and 41 equally abundant species, respectively. This observed drop in true diversity values (approximately 50%) in comparison to Shannon’s true diversity index is probably due to the unequal abundances in the data set where many OTUs can be considered rare by their frequencies and the greater influence of OTU abundances in Gini-Simpson’s index.

### Group significance as measured by beta diversity suggests that mangrove micro-habitats do not have distinctly different microbiomes

We used beta diversity distance metrics to assess the statistical significance of differences between the sample sites using the PERMANOVA pairwise non-parametric statistic test as shown in Supplemental Table 2. The results for the Bray-Curtis and Weighted Unifrac quantitative distance metrics (Fig 4) were used for the boxplots. These were selected since they account for both richness and abundance, and in order to analyze the data using both phylogenetic and non-phylogenetic approaches, thus providing a more complete information about the samples.

The results of the quantitative distance metrics indicate a heterogeneous data set with substantial variation both within and between groups. The corrected p-values for multiple tests (q-values) shown in the table 2 fails to reject the null hypothesis that different mangrove sites vary in their microbial composition (α= 0.05, 95% confidence). Defined as a geometric partitioning of multivariate variation in the space of a chosen dissimilarity measure (Anderson 2017), the pseudo-F test results for PERMANOVA are directly affected by the variability observed in the samples within groups, which is high and therefore may underlie the lack of significant differences and a weak effect of the group separation, despite the apparent dissimilarity between the groups in the plots and in the alpha diversity analyses where all the indices point to large differences between the sites. Notably, given the relative similarity of the micro-habitats, three replicates may not have enough statistical power to identify meaningful differences between sites using PERMANOVA.

### The influence of environmental variables

The environmental variables salinity, water content, organic matter and temperature from each study site were measured and associated to the community structure in the samples through Pearson’s correlation test and confirmed using a Mantel test. The results revealed a significant correlation between the prokaryotic communities and temperature and water content (Fig 5). The salinity and organic matter measures do not produce a distinguishable effect on the community structure in these sediment samples.

### Numerous taxa associated with nutrient cycling identified within communities

Analysis of microorganism taxonomies are often used in the literature to address the role that groups of microorganisms play in the environmental and geochemical aspects of nutrient cycling and metabolism, usually referring to specific transforming communities by the compounds they are associated with (Jørgensen, Findlay, and Pellerin 2019; Wasmund, Mußmann, and Loy 2017; Buan 2018; Bae et al. 2018; Li et al. 2015; Levipan et al. 2016). Here, we rely on previous works that have shown which taxa are the most probable drivers for some soil/sediment processes such as methanogenesis, nitrogen fixation and nitrification which are carried out by well-characterized microbial groups (Levipan et al. 2016; Fierer 2017) and use the information to link the families found in this study to their most probable and relevant nutrient cycling activities (Figure 6).

**Figure 6.**
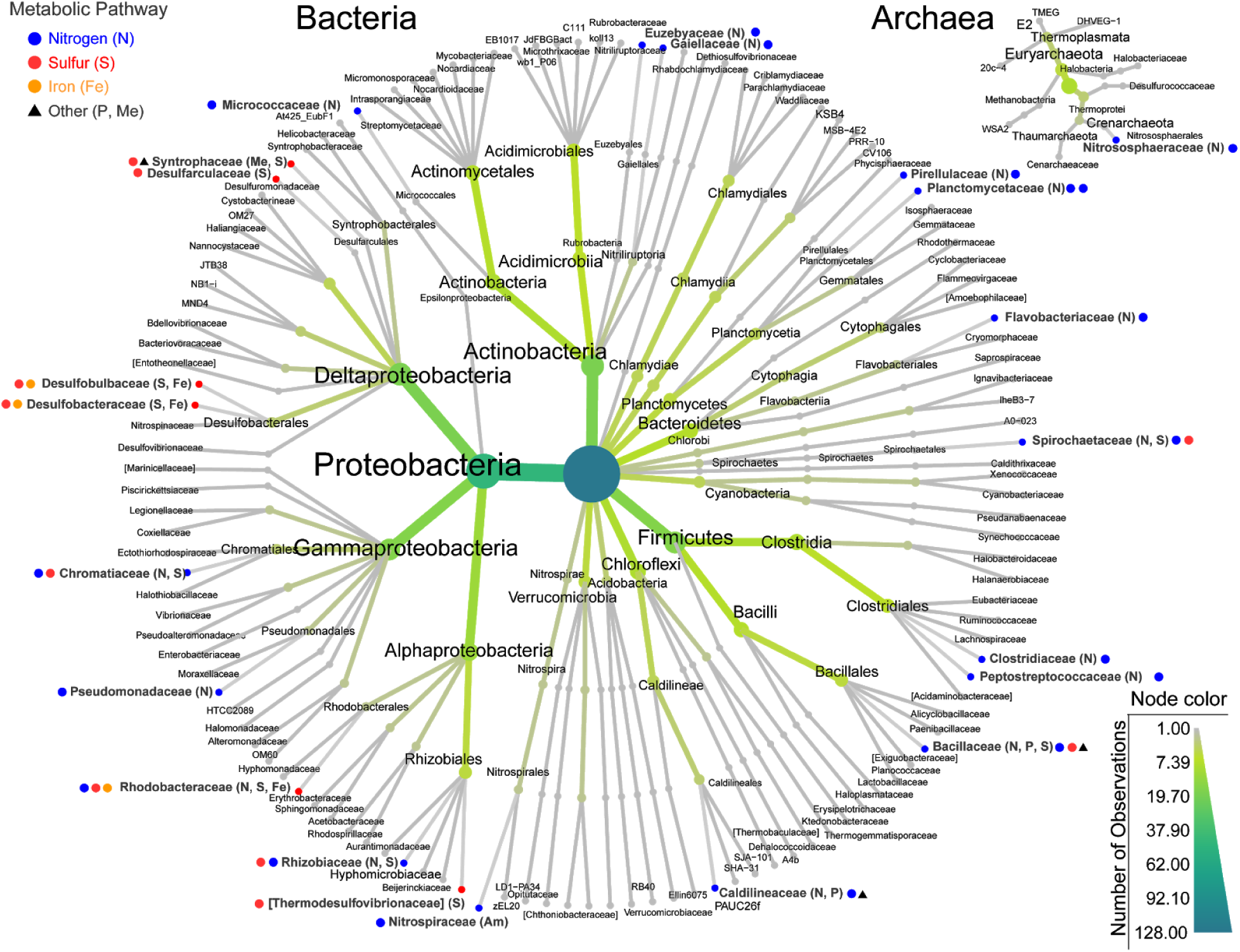
Phylogenetic tree showing additional metabolic data. The phylogenetic tree highlights the associations between the taxa and metabolic pathways that are possibly driven by the prokaryotic families in the sediments based on the literature.

The most abundant family observed in the samples was *Bacillaceae* (19,950 sequences) which is known to be one of the most robust bacterial groups and for participating in several important cycles in soil environments including carbon, nitrogen, sulfur, and phosphorous (Mandic-Mulec, Stefanic, and van Elsas 2015). Another abundant group found in the samples was *Clostridiaceae* (12,033 sequences) which have been reported to play roles on nitrogen-fixing (Wiegel, Tanner, and Rainey 2006) and is probably participating in this cycle on the mangrove sediments. The third most abundant family in the samples was *Hyphomicrobiaceae* (10,104 sequences) which is also possibly participating in the nitrogen-fixing and denitrification processes and likely has a role in sulfur cycling in the sediments (Wiegel, Tanner, and Rainey 2006; Oren and Xu 2014; Meyer et al. 2016). The anaerobic oxidation of ammonium (ANAMMOX) is considered as an alternative route for N loss to the denitrification process in environments with anoxic conditions and is driven by microorganisms that belong to the order *Planctomycetales* including the family *Planctomycetaceae* (Nie et al. 2015; Wang et al. 2012). The detection of *Planctomycetaceae* in the mangrove sediments indicate that they are the probable drivers for this part of the nitrogen cycle in this environment. The family *Caldilineaceae* is commonly found in studies concerning nitrogen removal in wastewater consortia and have also been correlated with phosphorus removal in wastewater treatment systems (Chen et al. 2019; Zhang, Xu, and Zhu 2017; Drury, Rosi-Marshall, and Kelly 2013) indicating the possibility that the group is active in N and P cycles in these sediments. The other families that present the capacity for nitrogen fixing in the sediments are *Flavobacteriaceae* (Kämpfer et al. 2015), *Pseudomonadaceae* (Özen and Ussery 2012) and *Spirochaetaceae* (Lilburn et al. 2001). The family *Chromatiaceae* has some representatives that are active in the nitrification process, as the genus *Nitrosococcus* (Campbell et al. 2011). Some representatives of the family *Gaiellaceae* are shown to reduce nitrate to nitrite (Albuquerque and da Costa 2014) similar to the family *Euzebyaceae* that is also capable of denitrification through the reduction of nitrate to nitrite and nitrite to N_2_ (nitrogen gas) (Kurahashi et al. 2010) completing the process of N loss. Organisms with reported capacity of performing the ammonification processes were also present, represented by the family *Peptostreptococcaceae* (Slobodkin 2014) and the family *Micrococcaceae* which has genes involved in ammonia assimilation (Dastager et al. 2014) as well as the family *Rhodobacteraceae* (Delmont et al. 2015). Ammonia-oxidizing organisms are also important to the environmental N cycle and can be divided in ammonia-oxidizing bacteria (AOB) which is represented in the samples by the family *Pirellulaceae* (Y.-F. Jiang et al. 2015) and ammonia-oxidizing archaea (AOA) which is represented by the family *Nitrososphaeraceae* (Kerou et al. 2016).

Some families in the sediments can be correlated to the sulfur cycle as is the case of the *Rhodobacteraceae* (Delmont et al. 2015). Another important family involved in the sulfur cycle was *Spirochaetaceae* which is reported to participate in the sulfur oxidizing process (Wiegel, Tanner, and Rainey 2006; Oren and Xu 2014; Meyer et al. 2016), while the sulfate-reducing step is possibly driven by organisms of the families *Thermodesulfovibrionaceae* (Bhatnagar et al. 2015; Wiegel, Tanner, and Rainey 2006; Oren and Xu 2014; Meyer et al. 2016) and *Desulfarculaceae* (Sun et al. 2010) comprising the sulfate-reducing bacteria (SRB) in the samples. *Chromatiaceae* is an important representative of a group known as purple sulfur bacteria and is strongly correlated with the sulfur cycle in the environment and in wastewater treatment plants (Wiegel, Tanner, and Rainey 2006; Oren and Xu 2014; Meyer et al. 2016; Hanada and Pierson 2006; Xia et al. 2019). Some members of the family *Syntrophaceae* are able to use sulfur or other sulfur compounds as electron acceptors, thus participating in the sulfur cycle in the environment (Kuever 2014) and are even capable of contributing in some steps of the methanogenesis by cooperation with methanogenic archaea (Cheng et al. 2013). The family *Desulfobacteraceae* has representatives that are capable of reducing iron and sulfate in the environment and are relatively abundant compared to other groups indicating that they might play an important role in these cycles in these mangrove sediments. The same environmental roles can be attributed to the family *Desulfobulbaceae* (Wiegel, Tanner, and Rainey 2006; Oren and Xu 2014; Meyer et al. 2016; Reyes et al. 2016). The other families that may be involved in the iron cycle in the sediments are *Flavobacteriaceae* that have several genes for iron acquisition and *Rhodobacteraceae* with the presence of genes that encode for its transport (Delmont et al. 2015).

### Microhabitats are differentially enriched in specific taxa

While the beta-diversity measure suggests that the different mangrove forest sites are not distinct ecologies, several lines of evidence, such as the significant environmental influence on community structure (Figure 5) and the distribution of taxa by site (Supplemental figures 2, 3 and 4) suggest that individual species may be enriched in specific sites. To evaluate this we tested each taxon at each site for significant differential abundances (see Methods). We found 10 taxa (3 orders, 4 families, 1 genera, and 2 species) that had significantly different abundances between sites (Fig 7, [Table 4]). Notably, five of these taxa (Fig. 7A-E) are also known to play important roles in nutrient pathways. We also observed that the Seco site had higher frequencies of observations (number of samples that observations occurred in) for 7 of these taxa, while the Submerged site only had 3.

**Figure 7.**
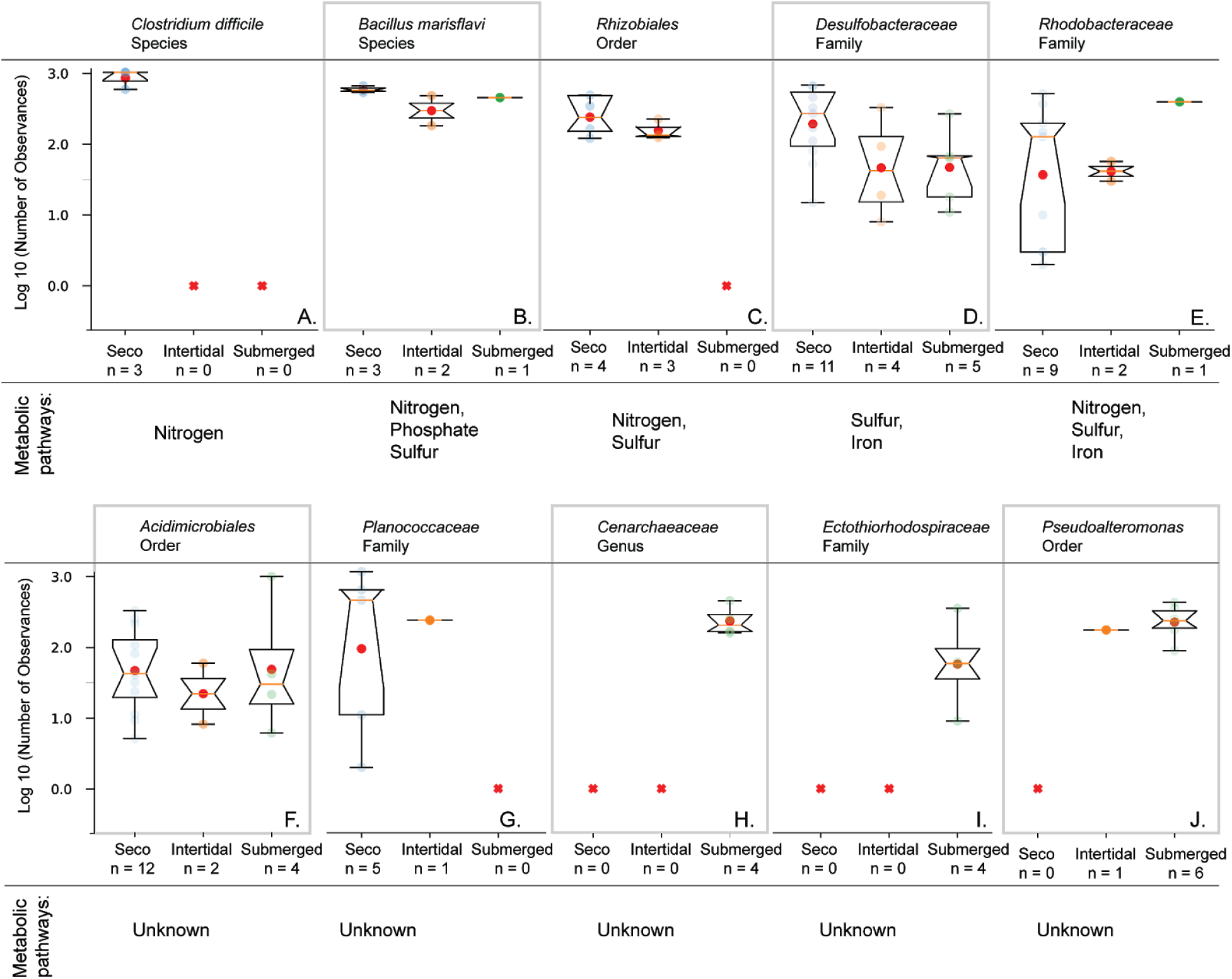
Site specific measures of taxa finds significant enrichment in Seco and Submerged sediments. To determine if individual taxa were differentially abundant at specific sites we looked for taxa that had significantly different abundances between sites (Chi-squared, p-val <= 0.05), statistically different distributions of abundance (Mann-Whitney U, p-val <=0.05), and whose mean effect size exceded 5% difference between all sites (see Methods). We found 10 taxa with significant enrichment at a particular site, 7 of these being more abundant at the Seco site (A-G) and 3 being highest at the Submerged site (H-J).

## DISCUSSION

The large number of taxa identified in the mangrove sediments in this study is in accordance with the related studies (Alzubaidy et al. 2016; Nogueira et al. 2015; Colares and Melo 2013; Andreote et al. 2012) in which the presence of many uncultured organisms that cannot be cultured using laboratory conditions were identified.

Previous work in mangrove microbial diversity found that composition of bacterial communities in sediments correlated with distinct hydrodynamic regimes, as well as characteristics like granulometry and organic matter content. Conversely, a study with mangrove sediments (Rocha et al. 2016) also observed the existence of different characteristics in the community structure of mangrove sites but found that it had greater dependence on vegetation, than the abiotic variables. Similarly, (Peixoto et al. 2011) observed bacterial profile clusters within the same mangrove, and suggested that abiotic factors or pollutant distributions can generate niche variations that could explain the differences. Using alpha-diversity tests we found that a greater number of OTUs, as well as a greater taxonomic diversity, are present in the submerged mangrove sediments, while the intertidal region has lower richness and diversity (Fig 3). We also performed a multivariate test on environmental variables and found that the absence or presence of water as well as temperature were important characteristics influencing the communities at the OTU level (Fig. 5) and, thus, could be important in explaining the variability observed in the alpha-diversity results. It is noteworthy that the two statistically relevant variables in this study present rapid changes in the mangrove areas, especially in the intertidal region and, thus, we hypothesise that the great variability in environmental parameters in this part of the mangrove restricts the options for some groups with specific necessities acting as a selective force eliminating all but the most resistant species, resulting in lower diversity indices for this site.

Although differentiation in mangrove sediment communities from sites with distinct biotic and abiotic characteristics has previously been reported (Rocha et al. 2016; Peixoto et al. 2011; X.-T. Jiang et al. 2013) these studies typically only test for alpha-diversity and environmental variables correlations, but do not perform a statistical analysis of the difference in communities between sites. Indeed, to our knowledge, the only group that has previously performed PERMANOVA analysis, (Ceccon et al. 2019) found that environmental factors have influence in the diversity between the sites, but no statistical differences were found in the group significance analysis, which is consistent with our findings. While the qualitative analysis of community diversity between mangrove sites suggests that micro-habitats have different microbial population structures these differences are not statistically significant likely as a result of extensive intrasite variation.

The Weighted Unifrac distance analyses (Fig 4; Supplemental Figure 6) takes into consideration the abundances and phylogenetic relations between the groups in the samples. Our results are in accordance with the premise that there exists a “core microbiome” in mangrove sediments that shows the dominance of some specific taxa (Cabral et al. 2016; Andreote et al. 2012; Mendes and Tsai 2014; Imchen et al. 2018; Rocha et al. 2016; Peixoto et al. 2011) suggesting that environmental factors correlated to microbiome variations only account for differences in the lower taxonomic levels and that the phylogenetic correlations and abundances in these ecosystems are relatively stable.

Part of the recovered families in the analyses could not be assigned to specific functional roles and some of them are even still not officially assigned to upper taxa levels, which may reflect characteristics of sediments microbiomes that show enormous diversity and many organisms that cannot be studied in laboratory conditions. From the families that could be assigned to a taxonomy, some were associated with specific roles in nutrient cycling (Fig 6). This motivated us to ask if specific sites had significant differential abundances of taxa. We found 10 taxa (3 orders, 4 families, 1 genera, and 2 species) had significantly different abundances between sites (Fig 7), five of which we had previously found to be associated with nutrient cycling. This suggests that, although the sites may not be distinct ecosystems (having failed to reject the null using beta-diversity tests), they nonetheless harbor different species in different abundances which are associated with different functional roles.

The identification of families associated with the diverse nutrient cycles in mangrove sediments was expected due to the previous observations of the astonishing microbial diversity and geochemical importance of prokaryotes in these environments (Singh et al. 2005; Andreote et al. 2012; Imchen et al. 2018); Zhao and Bajic 2015) and confirmed in this study. Along with the taxonomic and functional diversities observed in the organisms it was possible to identify that the different taxa in the data set presented diverse optimum oxygen conditions being some strict aerobic or anaerobic organisms as well as facultative anaerobic which can be correlated to the existence of oxic and anoxic zones in the superficial parts of the sediments in the mangroves (Singh et al. 2005).

Our study suggests that while the microhabitats have relatively large biotic and abiotic differences, including some taxa that have significantly different abundances, in general they have relatively similar population structures. One way to interpret this is to consider that both the seco and submerged sites may be acting as species reservoirs for the larger habitat, dispersing species through tides, storms, and other disruptive environmental events. Coupled to this is the conceptualization of the intertidal region as a hybrid zone with very strong selective pressures, thus acting as a barrier to many of the species present in each reservoir. Furthermore, while they may be abiotically different they all possess a common foundational species (Angelini et al. 2011), mangrove plant itself. It is worth considering that the microhabitats must act in concert to keep the mangroves themselves alive and as such there may be higher orders of ecological structure at play beyond, and in between, the boundaries of the microhabitats themselves.

## CONCLUSIONS

Our study found a large amount of microbial biodiversity in mangrove sediments, with distinct regions (submerged, intertidal, seco) exhibiting variance in species diversity as calculated by alpha diversity and taxa enrichment in some sites compared to the others, potentially due to differences in environmental variables that showed significant influence in the communities. While differences in microbial communities were observed between regions, these trends are not statistically significant. The taxonomy analyses show the dominance of some groups in the upper taxonomic levels in agreement with previous studies and the phylogenetic measures of the site samples suggest that the microbial communities are broadly similar. We identified specific families that could play a functional role in the geochemical cycles of different nutrients in this ecosystem. Finally, we observed that the intertidal region, the most biophysically dynamic site in the mangrove forest, presents the least biodiversity. That a dynamic cyclic environment could prove to be more harsh and selective niche is worthy of further study. Further exploration of this interesting point and confirmation of organisms groups that actively participate in element cycling and the corresponding transformation rates is the subject of future studies.

## ACKNOWLEDGEMENTS

The authors thank the Center for Genomics and Bioinformatics (CeGenBio) of the Federal University of Ceará (UFC)- Brazil, for the sequencing of samples. This study was financed in part by the Coordenação de Aperfeiçoamento de Pessoal de Nível Superior – Brasil (CAPES) – Finance Code 001.

## SUPPLEMENTAL FIGURES

**Supplemental Figure 1.**
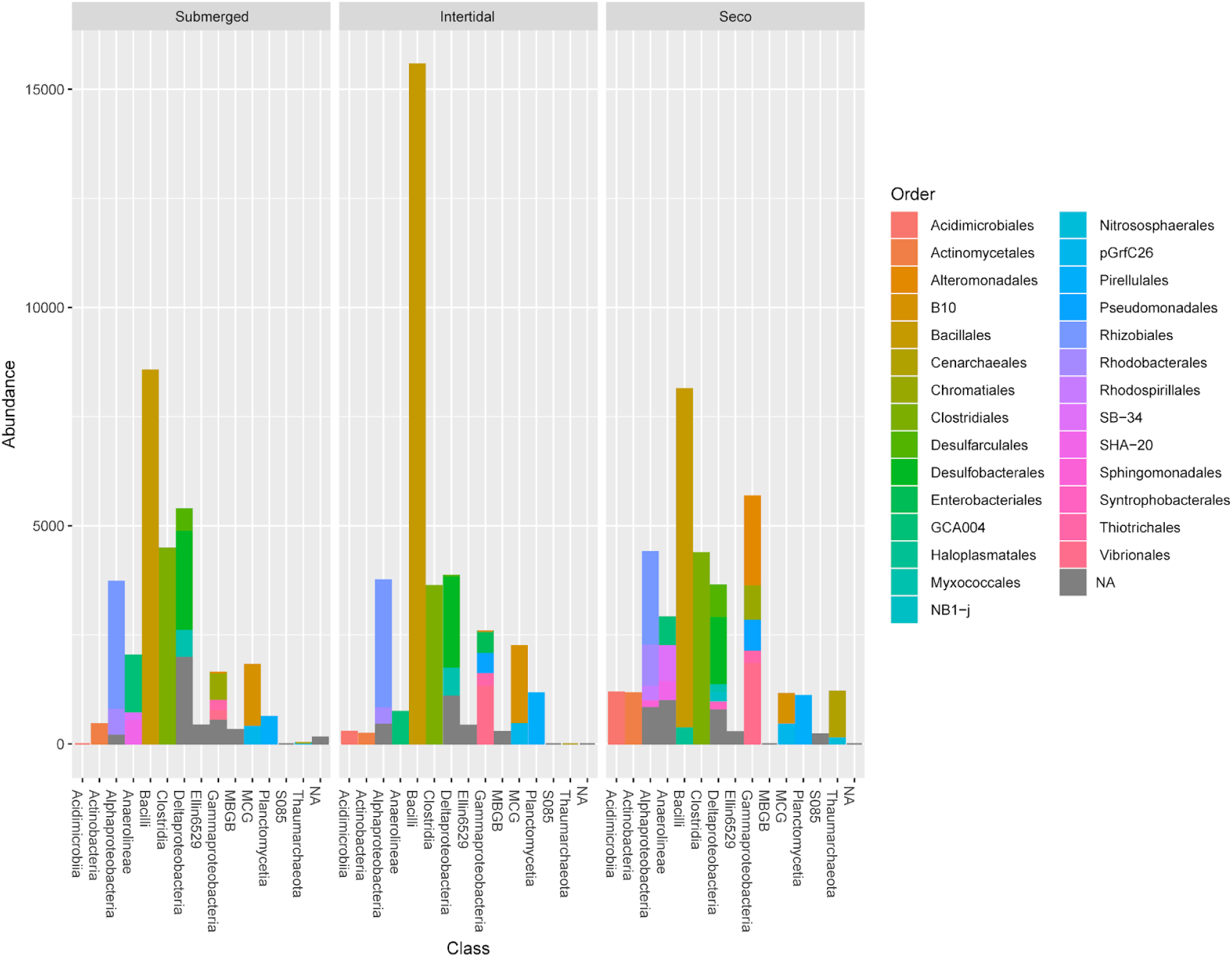
Abundance of Class and Order by site.

**Supplemental Figure 2.**
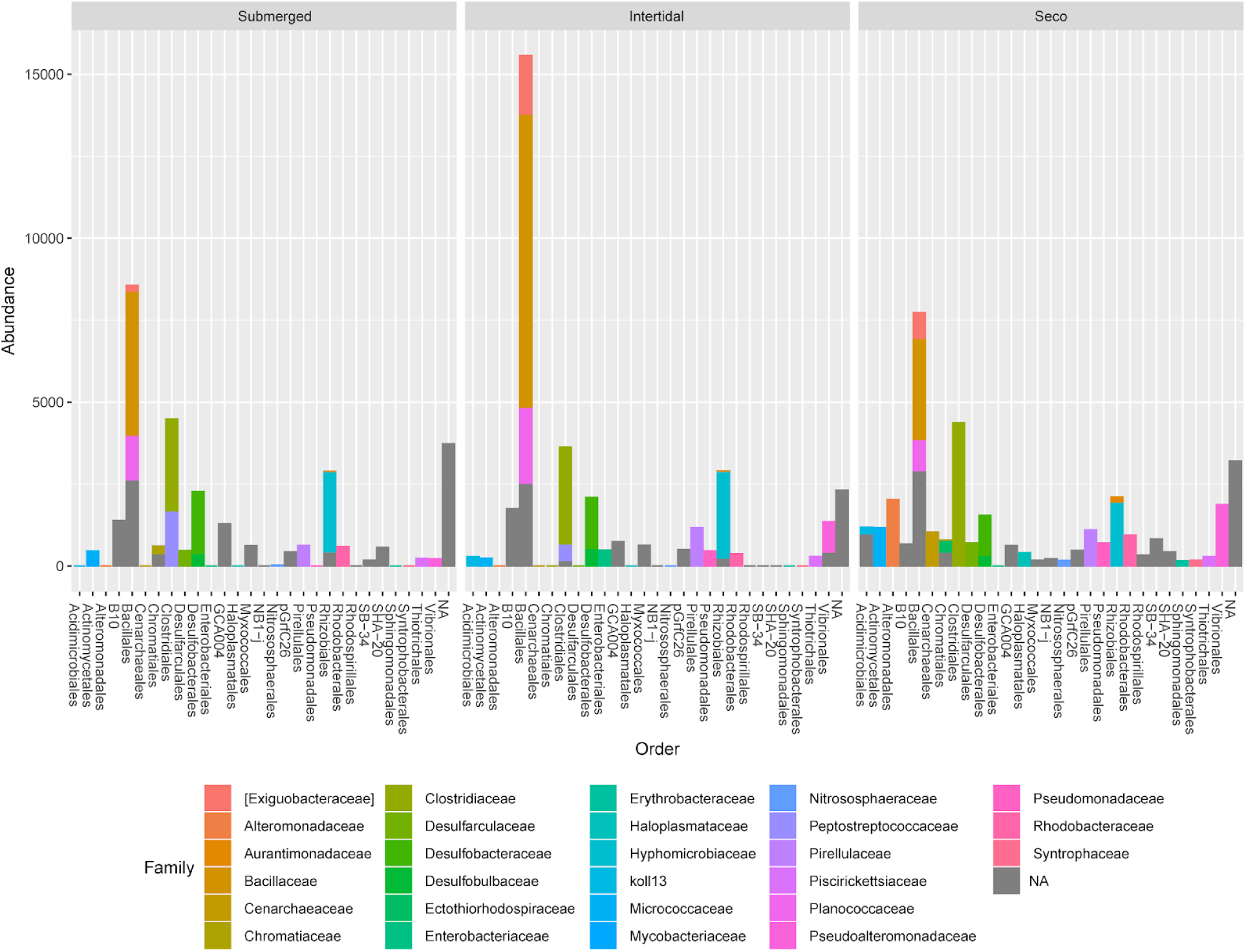
Abundance of Order and Family by site.

**Supplemental Figure 3.**
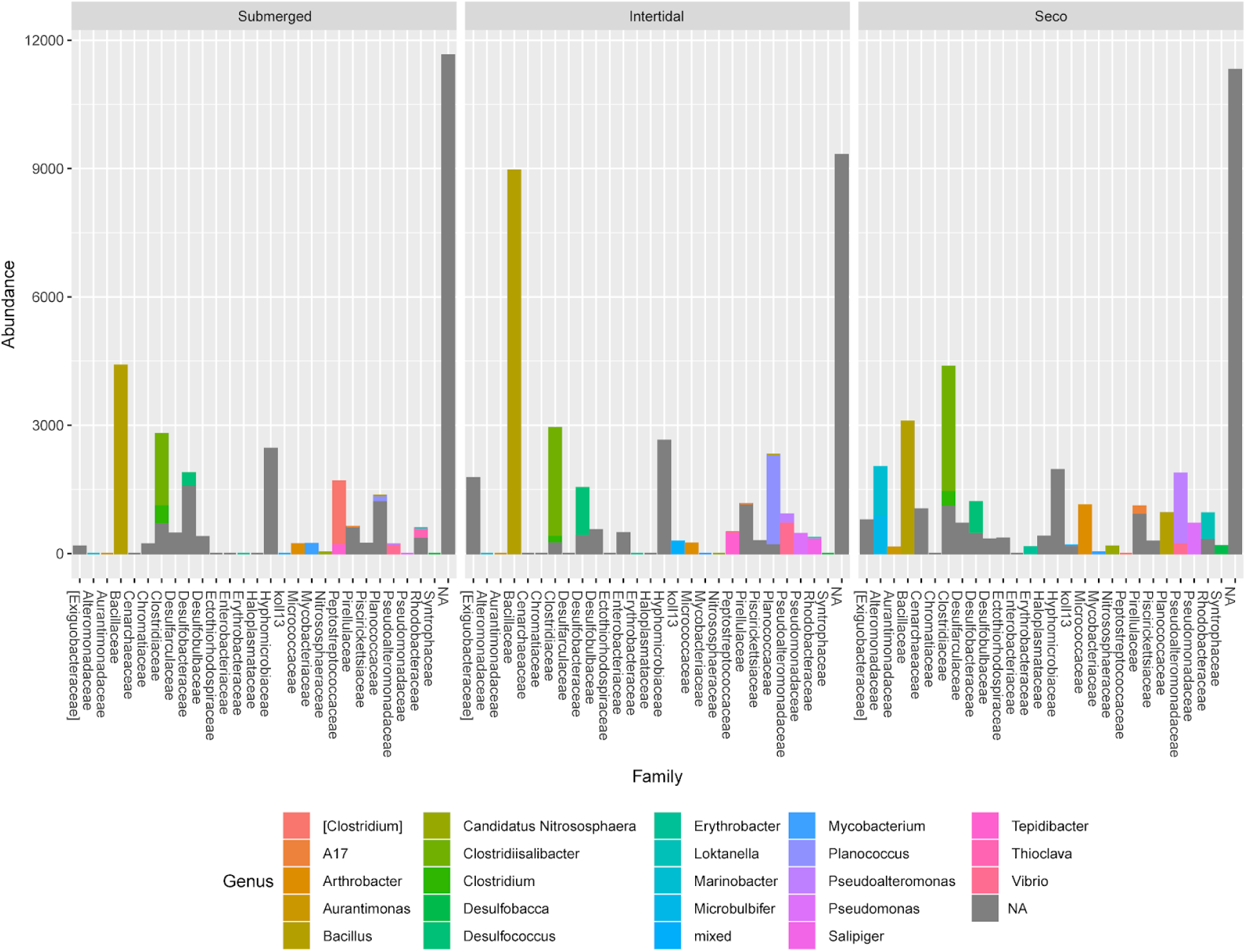
Abundance of Family and Genus by site.

**Supplemental Figure 4.**
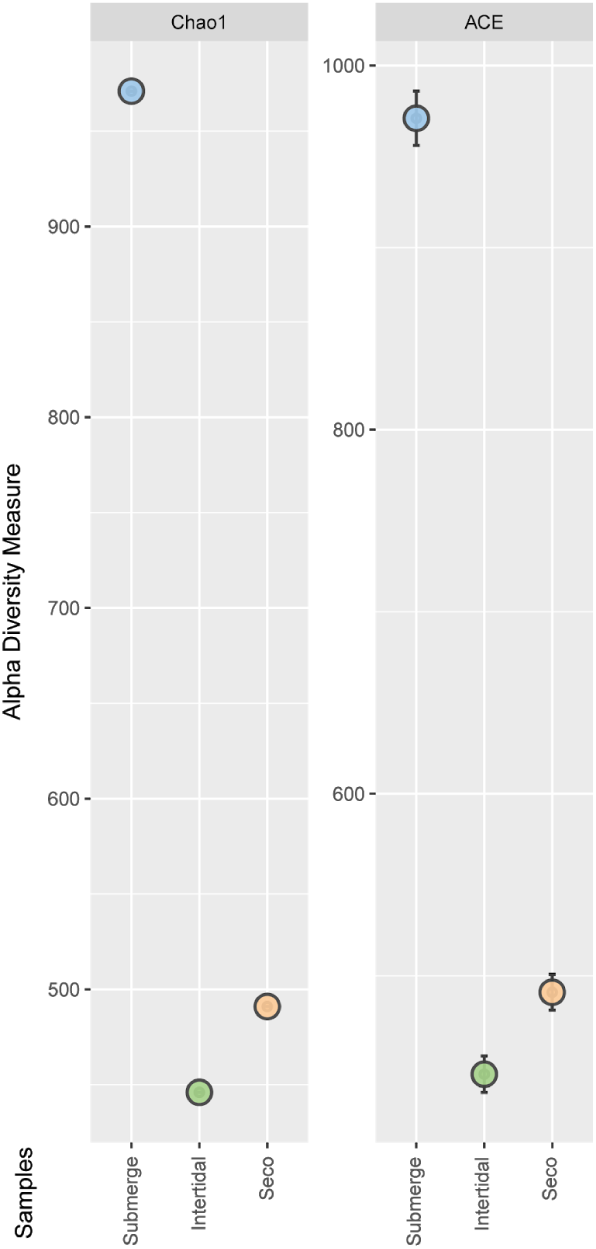
Additional Alpha diversity measures. In addition to the Alpha-diversity metrics described above we also used additional estimators. The Chao1 estimator calculates the estimated true richness of a sample and shows the same pattern observed for OTU richness, which is in agreement with the observation that the sequencing depth was enough to assess all the diversity in the samples. Based on the sample coverage, the ACE (Abundance-based Coverage Estimator) method accounts for possible missing taxa and estimated very similar OTU richness as the previous measures.

**Supplemental Figure 5.**
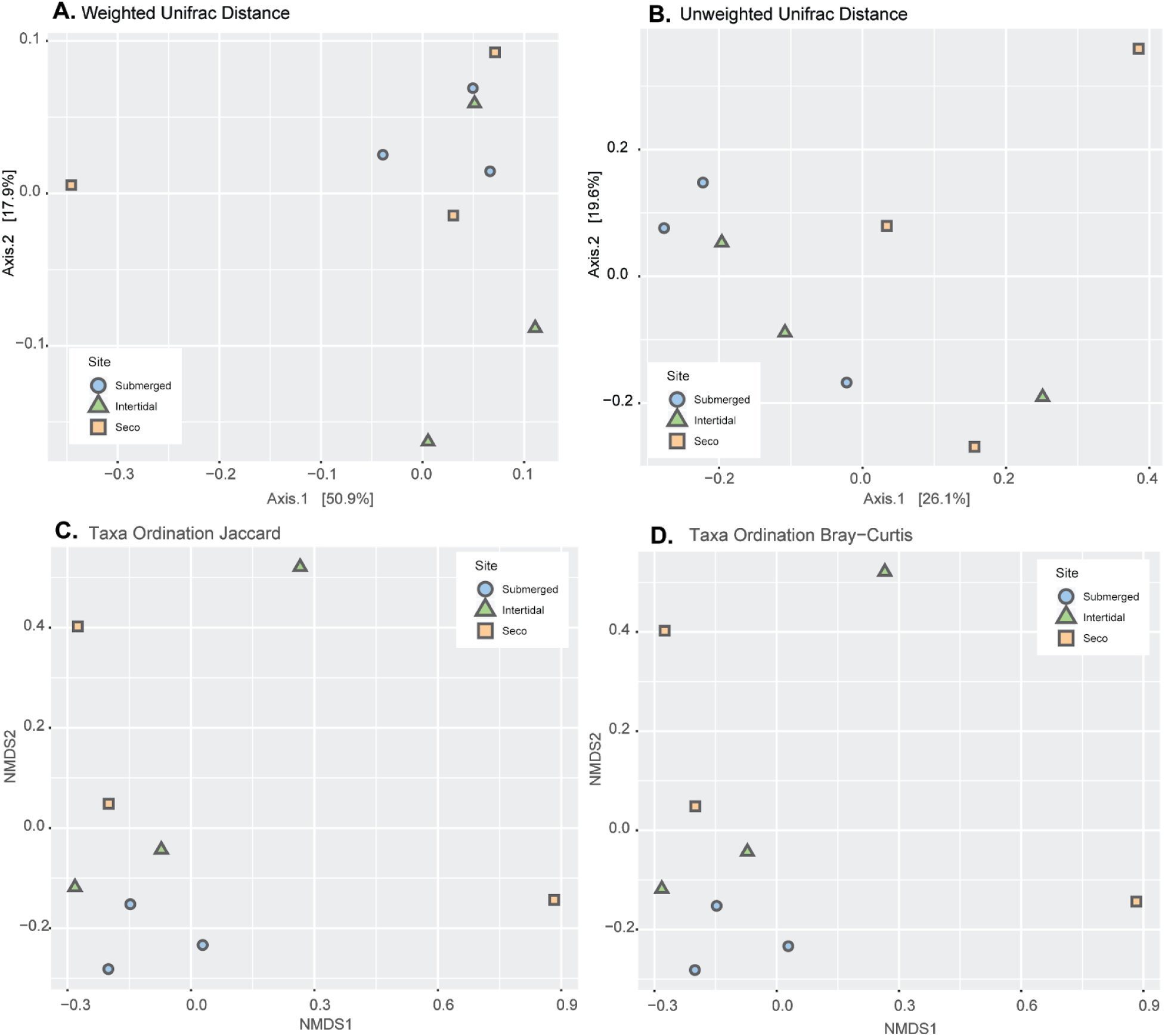
Non-metric distance scale (NMDS) analyses were used to plot the variability of samples within and between sites obtained by the non-phylogenetic distance metric methods Jaccard and Bray-Curtis. The Jaccard distance metric plot displays qualitative variability in the samples, meaning that the distance is based on the presence/absence of OTUs in each analyzed sample. Considering the observed OTUs per sample, we do not see a consistent clustering among the sites, suggesting that the differences in the OTUs found in each sample are not explained by the mangrove area where the sediment was collected. The Bray-Curtis distance plot shows the quantitative variability observed in the data, which accounts for the OTUs abundance in the samples. The diversity in the samples produced a weak clustering for the submerged site group indicating that the samples from this site were more homogeneous in terms of OTU abundances than the other two sites. Furthermore, we identify that the OTU abundance observed in the seco site samples was more heterogeneous than the other sites and also was a greater distance from the other two mangrove sites. Principal Coordinates Analysis (PCoA) plots were used to assess the phylogenetic distance metrics. The qualitative Unweighted Unifrac distance analysis measures the variability between groups of samples by assessing the correlations between the taxa observed in samples from specific sites. The PCoA plot shows that the samples do not cluster by site but also don’t present great differences in the phylogenetic correlations of the microbial taxa. In a quantitative approach the Weighted Unifrac distance metric uses the abundance information to compare the samples and assess distances within and between groups. When comparing samples by abundance of phylogenetically correlated taxa all three sites show similar results with most samples closely positioned in the plot. Both Unweighted and Weighted Unifrac found the seco site samples to be the greatest distance from any group while the submerged site was found to have have the smallest distances. All the distance metrics used for comparing the prokaryotic communities in the three mangrove site sediments show that it is not possible to clearly differentiate between the sites by the composition of microorganisms. While the submerged group samples showed a greater tendency to cluster compared to the other two site samples, this was not statistically robust given the composition of all sites.

**Supplemental Figure 6.**
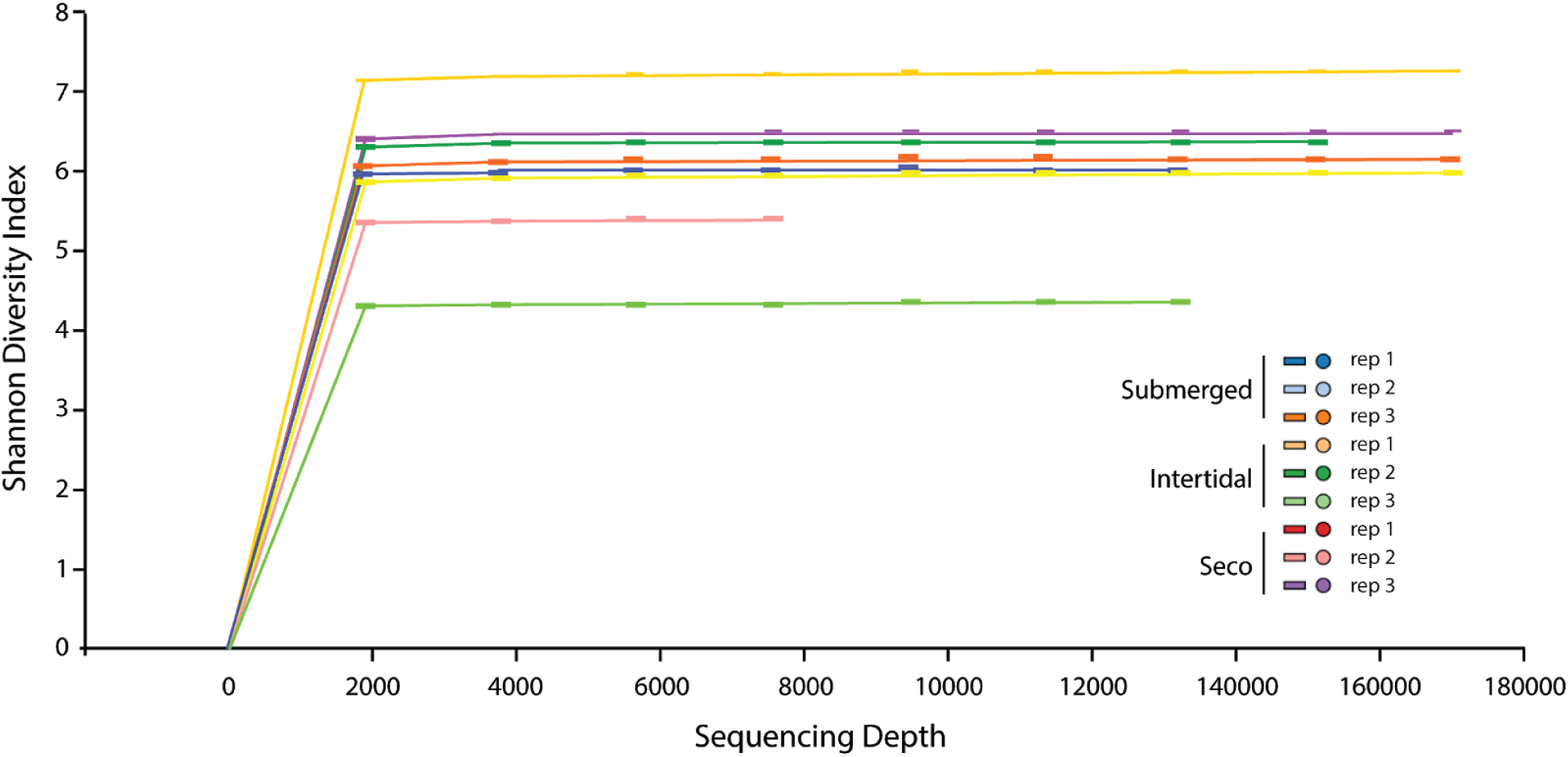
Alpha-rarefaction plot. An alpha rarefaction curve plot was generated in order to verify the efficiency of the sequencing depth used in the study and it shows that the number of observations was enough to assess all the diversity in the samples Here we show the Shannon Diversity Index of random sub-samples of the sequence libraries from each replicate of each site. In each case we find the diversity is effectively saturated at a sequencing depth of 4000 suggesting that each replicate was sufficiently sequenced..

## SUPPLEMENTAL TABLES

**Table 1.** Frequency and abundance of the observed taxa at the phylum level.

| Phylum | Sequences | % Sequences | OTUs | % OTUs |
| --- | --- | --- | --- | --- |
| [Caldithrix] | 109 | 0.053 | 6 | 0.19 |
| [Parvarchaeota] | 39 | 0.019 | 7 | 0.23 |
| AC1 | 36 | 0.017 | 6 | 0.39 |
| Acidobacteria | 7524 | 3.72 | 78 | 5.00 |
| Actinobacteria | 10853 | 5.366 | 99 | 6.04 |
| Armatimonadetes | 14 | 0.006 | 1 | 0.06 |
| Bacteroidetes | 5918 | 2.926 | 7 | 0.40 |
| BRC1 | 31 | 0.015 | 84 | 4.81 |
| Caldithrix | 678 | 0.335 | 8 | 0.44 |
| Chlamydiae | 120 | 0.059 | 18 | 0.98 |
| Chlorobi | 359 | 0.177 | 7 | 0.38 |
| Chloroflexi | 17629 | 8.716 | 163 | 8.75 |
| Crenarchaeota | 10964 | 5.418 | 57 | 2.81 |
| Cyanobacteria | 1467 | 0.725 | 27 | 1.30 |
| Euryarchaeota | 304 | 0.15 | 17 | 0.81 |
| Firmicutes | 60618 | 29.971 | 116 | 5.45 |
| Gemmatimonadetes | 2062 | 1.019 | 6 | 0.27 |
| GN02 | 20 | 0.009 | 9 | 0.40 |
| GN04 | 107 | 0.052 | 25 | 1.11 |
| Hyd24-12 | 10 | 0.005 | 1 | 0.04 |
| KSB3 | 35 | 0.017 | 2 | 0.09 |
| LD1 | 6 | 0.003 | 2 | 0.09 |
| NC10 | 35 | 0.017 | 1 | 0.04 |
| Nitrospirae | 2086 | 1.031 | 3 | 0.13 |
| NKB19 | 65 | 0.032 | 12 | 0.52 |
| OD1 | 38 | 0.018 | 13 | 0.56 |
| OP11 | 10 | 0.005 | 2 | 0.09 |
| OP3 | 6 | 0.003 | 1 | 0.04 |
| OP8 | 3812 | 1.884 | 19 | 0.82 |
| OP9 | 7 | 0.003 | 2 | 0.09 |
| Planctomycetes | 11592 | 5.731 | 227 | 9.70 |
| Proteobacteria | 61889 | 30.599 | 391 | 15.23 |
| SBR1093 | 36 | 0.017 | 2 | 0.07 |
| SC4 | 65 | 0.032 | 5 | 0.17 |
| Spirochaetes | 431 | 0.213 | 12 | 0.40 |
| Synergistetes | 4 | 0.002 | 1 | 0.03 |
| Tenericutes | 17 | 0.008 | 14 | 0.47 |
| TM6 | 195 | 0.096 | 1 | 0.03 |
| TM7 | 2 | 0.001 | 2 | 0.07 |
| Verrucomicrobia | 576 | 0.26 | 37 | 1.23 |
| WS2 | 26 | 0.012 | 3 | 0.10 |
| WS3 | 2458 | 1.215 | 61 | 2.01 |
| TOTAL | 202253 | 99.957 | 1555 | 100.00 |

**Table 2.** Statistical significances (pseudo-F tests) for PERMANOVA beta-diversity analysis.

| Compared sites | Bray-Curtis |  |  | Jaccard |  |  | Weighted Unifrac |  |  | Unweighted Unifrac |  |  |
| --- | --- | --- | --- | --- | --- | --- | --- | --- | --- | --- | --- | --- |
|  | Pseudo-F | p-value | q-value | Pseudo-F | p-value | q-value | Pseudo-F | p-value | q-value | Pseudo-F | p-value | q-value |
| Submerged to Intertidal | 1.153651 | 0.209 | 0.3135 | 1.026788 | 0.201 | 0.3015 | 1.045695 | 0.642 | 0.642 | 1.088375 | 0.279 | 0.4185 |
| Submerged to Seco | 1.341529 | 0.103 | 0.3090 | 1.070966 | 0.086 | 0.2580 | 1.269203 | 0.178 | 0.444 | 1.149417 | 0.212 | 0.4185 |
| Intertidal to Seco | 0.911451 | 0.808 | 0.8080 | 1.013168 | 0.38 | 0.3800 | 1.206730 | 0.296 | 0.444 | 0.875849 | 0.786 | 0.7860 |
| all groups | 1.12715 | 0.116 | - | 1.03694 | 0.056 | - | 1.19704 | 0.162 | - | 1.03196 | 0.294 | - |

**Table 3.** Environmental variables measured at the sampling sites.

| Sample | Water content | Salinity | Temperature (celsius) | Organic Matter (%) |
| --- | --- | --- | --- | --- |
| P1 | submerged | 665,9 | 25 | 3.71 |
| P2 | intertidal | 677,6 | 29 | 3.86 |
| P3 | dry | 1.047 | 29 | 1.4 |

**Table 4.**
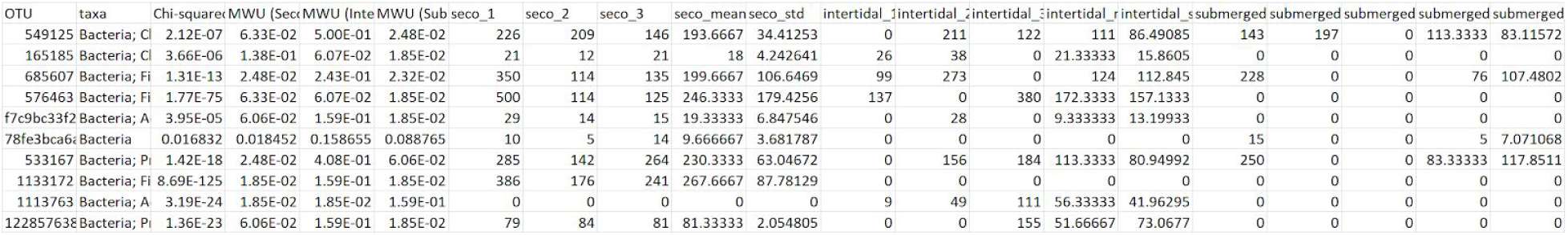
Site Specific OTUs.

